# Rnalys: An Interactive Web Application for Transcriptomic Data Analysis and Visualization

**DOI:** 10.1101/2025.02.12.637847

**Authors:** Christoffer Frisk

## Abstract

**Background:** The rapid growth of biological data from omics and transcriptomics research necessitates tools that streamline data visualisation and interpretation. Existing solutions for RNA-Seq analysis are often complex. Rnalys was developed as a lightweight, open-source web application tailored for the downstream analysis of counted and quality-controlled reads. It is designed as a tool to identify outliers and efficiently handle datasets with multiple batches, tissues, and disease types. Additionally, Rnalys provides representation of biological relationships, enabling users to perform comprehensive exploratory data analysis and generate publication-quality figures.

**Results:** Rnalys leverages Plotly’s Dash framework to provide an intuitive, interactive interface for the exploration and analysis of high-dimensional genomic data. Users can perform a range of analyses, including sample selection, batch correction, data transformation, gene expression exploration, PCA, differential expression analysis, and gene set enrichment analysis. The platform offers dynamic parameter adjustments and real-time display, enhancing data exploration and analysis.

**Conclusion:** Rnalys simplifies the process of data handling and visualization, allowing researchers to concentrate on interpreting biological results. The open-source nature of the tool, coupled with its user-friendly interface, facilitates customization and integration into existing workflows, making it a valuable resource for both individual researchers and collaborative teams.

## Background

The rapid growth of biological data from omics and transcriptomics research necessitates tools that streamline data visualization and interpretation. Existing solutions for RNA-Seq analysis, such as Galaxy [1, 2], cover comprehensive workflows but can be complex and resource-intensive for users focused solely on downstream analysis. To address this, Rnalys was developed as a lightweight, open-source web application tailored for the downstream analysis of counted and quality-controlled reads.

This application provides a range of data visualization and exploration tools for bioinformatics analysis. Its interactive, easy-to-use interface helps both new and experienced researchers quickly and effectively gain insights from their data. The application is open-source under the MIT licence, with the full code base available on GitHub, allowing customization to meet specific research needs, such as adding functions for differential expression (DE) analysis and making necessary modifications.

## Implementation

Rnalys is developed primarily in Python (v3.7+) using Plotly Dash [3] for the front-end interface, with R (v4.1.2) scripts supporting backend analyses via DESeq2 [4], edgeR [5] and limma [6] libraries. The application runs locally, either on individual machines or an internal network, with the optional feature of Enrichr analysis requiring external connection. Analyses are stored continuously for quick retrieval, enhancing workflow efficiency.

### Data Upload and Setup

The platform consists of a data upload/setup page and a visualisation/exploration area. Users upload a count file of raw gene expression counts and a metadata file specifying experimental conditions (Figure 1). The interface supports drag-and-drop or file selection, with successful uploads visually confirmed. Users can select two variables for analysis and one batch variable for subsequent visualization. The user is also given the option to select an example data set which generates a two-tissue-based dataset of negative binominal count data.

**Figure 1.**
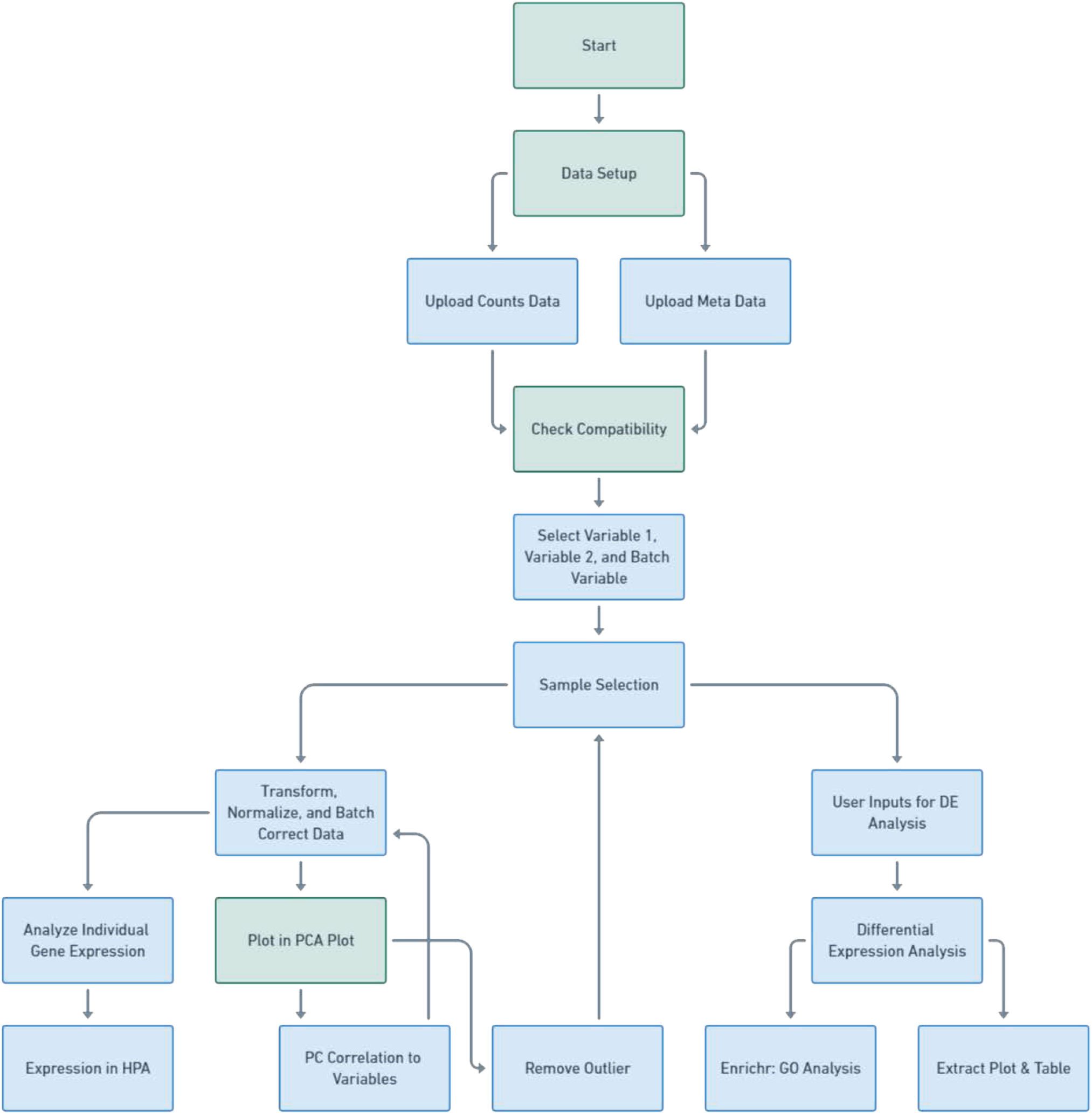
RNA Analysis Flowchart. The flowchart details the series of steps in the data analysis procedure, beginning with ‘Data Setup’ and moving through stages such as ‘Upload Counts Data’, ‘Plot in PCA Plot’, and ‘Differential Expression Analysis’. Steps that require user input are emphasized in blue.

### Data Selection and Visualization

During the sample selection process, data can be filtered by selecting specific samples based on chosen variables and batch factors from the initial page. There is also an option to exclude certain samples from the analysis. For data transformation and normalization, options include variance stabilization, r-log transformation, and mean of ratios normalization, all accessible via a drop down menu. These transformations use built-in functions from DESeq2. Additionally, confounding variables can be removed using the limma R-package. Once submitted, the calculations are run in the background, and the results populate the subsequent plots during the exploratory stage. Data selections can be saved and reloaded the next time the application is started. Count data by gene can be displayed through box plots grouped by the selected variable, and named by Ensembl or HGNC ID. For datasets with Ensembl IDs, there is also an option to search for the expression of genes in various human tissues according to the Human Protein Atlas [7] (Figure 2a).

**Figure 2.**
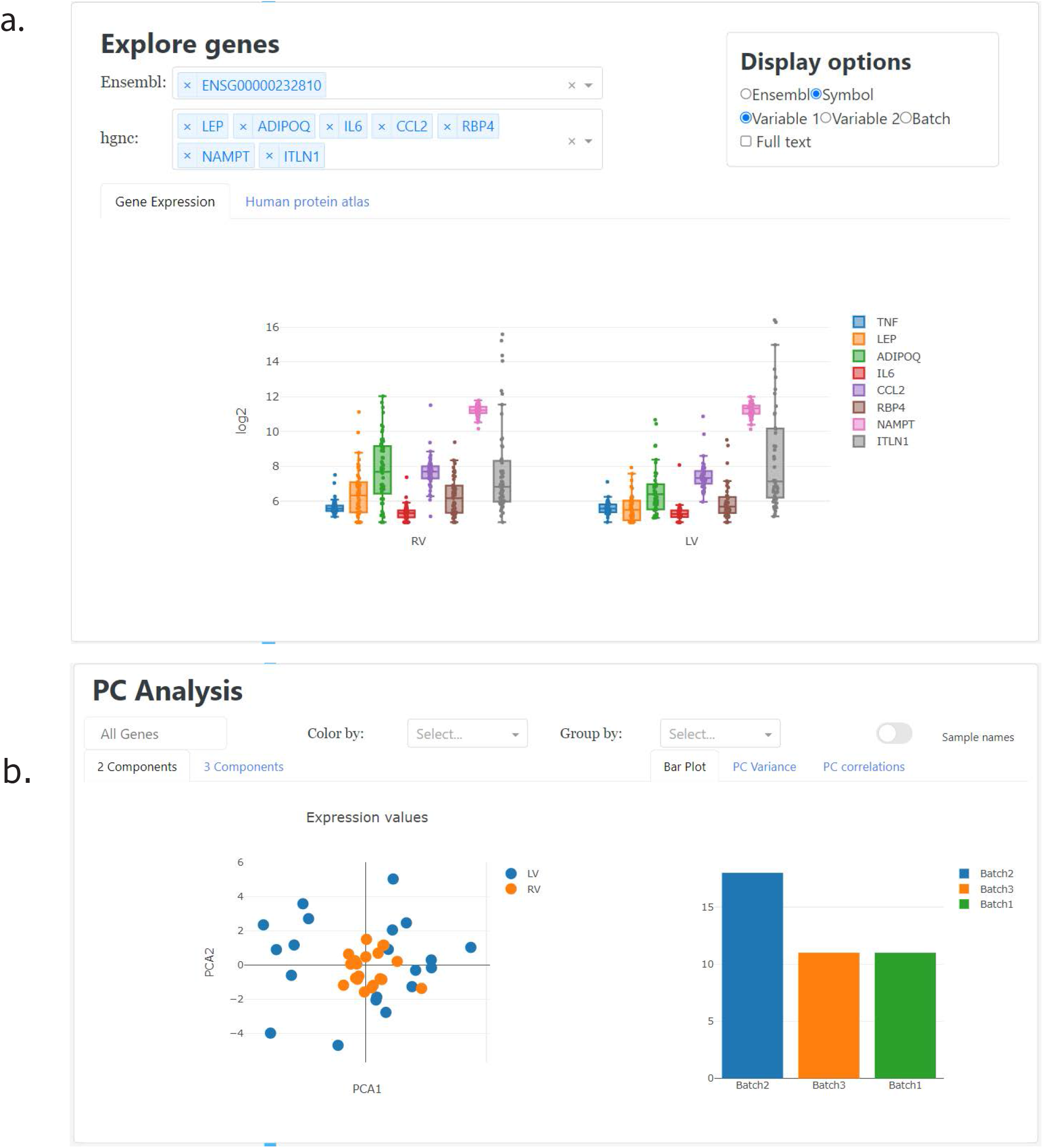
(**a)** The user interface for gene exploration enables users to select genes from the dataset using either Ensembl IDs or HGNC symbols. The chosen genes are grouped based on variables selected from the “Display Options” menu and visualized in box plots. Users can hover over the plots to view the specific sample names represented. (**b)** This figure provides an example of a dataset comparison between two tissue types. The left panel shows a principal component (PC) plot of the first two components, while the right bar plot displays the number of samples for each tissue type, as specified by the “Group by” dropdown menu.

A principal component analysis (PCA) is performed on the selected transformed or normalized data, with the default setting analysing all genes. However, users have the flexibility to adjust the number of genes included in the analysis for exploratory purposes. The PCA results are visualised in scatter plots displaying either two or three components, providing insights into the underlying data structure (Figure 2b). Additionally, sample distribution based on batch, tissue, or other metadata variables is depicted in bar plots, alongside visualizations of PC variance and correlations with metadata.

Differential expression analysis can be conducted using either DESeq2 or edgeR, with options available for low gene count filtering and the specification of a design formula and reference. The analysis is run on the raw count data with new parameters and not on the results from earlier in the analysis. The results are presented in an interactive volcano plot generated using the dash-bio library, allowing users to hover over points to highlight specific genes of interest (Figure 3a). When employing DESeq2, an “MA-plot”, is also available, where the y-axis represents the log2 fold-change and the x-axis shows the mean of normalized counts. The volcano plot includes interactive features such as a range slider for gene highlighting and the ability to set an adjusted p-value threshold for identifying differentially expressed genes, which can be exported as a table.

**Figure 3.**
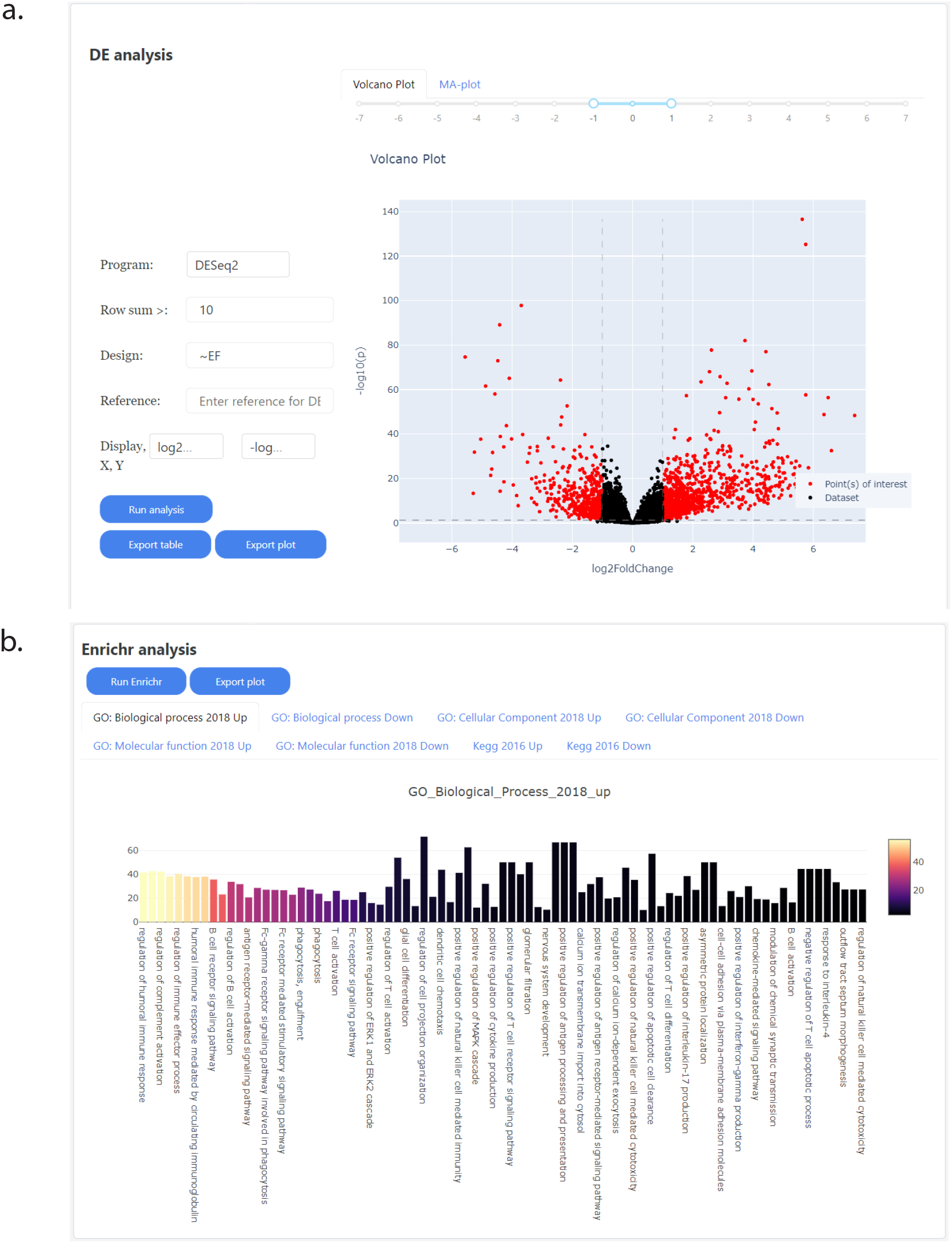
**(a)** This figure shows a DESeq2 analysis where genes with a total count below ten have been filtered out. The design formula is based on tissue type. Users can hover over the scatter plot to view details about highlighted genes. Additionally, the fold change in relation to the mean normalized counts can be examined in the accompanying MA plot. **(b)** This panel presents an example of an Enrichr analysis, displaying the enrichment of down-regulated genes within Gene Ontology (GO) biological processes. The y-axis represents the percentage of genes involved in each GO term, with colours indicating the adjusted p-value: lighter colours represent higher p-values, while darker colours represent lower p-values.

Furthermore, these visualizations can be exported as publication-ready plots using R scripts that reference the required files. The scripts, once executed, generate customizable volcano plots using the EnhancedVolcano package [8], allowing for customization to specific needs.

### Enrichment analysis

The final tool is an enrichment analysis which utilizes Enrichr [9] on the selected DE genes from the volcano-plot, separating them into up- or down-regulated. Tabs are provided for viewing the enrichment results from databases, which include enrichment in databases such as the Gene Ontology—covering biological processes, cellular components, and molecular functions—and the Kyoto Encyclopedia of Genes and Genomes (KEGG) (Figure 3b). There is an option to export the data and generate a plot script, which allows for customization and plots better suited for publication.

### Data Storage and Session Tracking

When a new set of samples and a specific transformation or normalization is selected, a new session is initiated, identified by a unique seven-character session ID. This ID is used to create a new subset of count files and metadata files, which are stored in a folder named ‘data/generated’. The details of this process are documented in a text file called “session_file.txt” and saved under “data/datasets”.

Additionally, the selected sample set can be saved for reloading after the application has been closed. This information is stored in the file “datasets.csv” under “data/datasets”. The scripts for DESeq2, edgeR, and limma are located in the ‘/functions’ folder. Furthermore, scripts for creating volcano plots are generated from a template stored in the “data/templates” folder and are saved in “data/scripts”, with the file name corresponding to the session ID.

## Results and Discussion

Rnalys enhances data processing through an intuitive interface that supports differential expression analysis, principal component analysis (PCA), and enrichment analyses with dynamic visualizations using Plotly’s Dash. Designed specifically for the downstream analysis of counted and quality-controlled reads, it provides an interactive platform for data visualization and exploration, allowing users to focus on interpreting their data.

PCA and correlation visualizations help identify patterns, outliers, and the influence of variables on gene expression. Users can conduct interactive differential expression analysis using DESeq2 or edgeR, with exportable plots and R scripts available for further customization and creation of publication-quality figures. An example dataset featuring two tissue types with negative binomial count data is included to help users familiarize themselves with the tool’s functionalities before analysing their own data.

Optimized for handling large transcriptomic datasets, Rnalys is equipped to manage complex experimental designs involving multiple batches, tissues, and conditions. It includes features for batch correction and outlier detection, which are important for managing high-dimensional datasets. The application speeds up analysis by saving and reloading previous calculations, with the option to force recalculations as needed.

While several software tools are available for RNA-Seq data analysis, each offering functionalities for different stages of the workflow, Rnalys focuses on simplifying the downstream stage for users who have already prepared their data. Unlike platforms such as Galaxy [1, 2], which cover parts of what Rnalys is providing, Rnalys offers interactive plots and lightweight solution optimized for handling high-dimensional biological data.

Future development includes bridging the gap between processing steps in the application and manual work on the generated data. This includes generating a script tracking the commands leading up to the different outputs of the analysis.

## Conclusion

Rnalys offers an interactive platform for visualizing and exploring high-throughput biological datasets, integrating differential expression and enrichment analysis to uncover underlying biological processes. Its customizable, user-friendly design makes it suitable for both individual researchers and collaborative teams.

## Availability and requirements

**Project name:** Rnalys

**Project home page:** https://github.com/Christofferfrisk/rnalys

**Operating system(s):** Platform independent

**Programming language:** Python, R

**Other requirements:** Python 3.7 or higher, R 4.1.2 or higher

**License:** MIT license

**Any restrictions to use by non-academics:** None

## Declarations

### Ethics approval and consent to participate

Not applicable

### Consent for publication

Not applicable

### Availability

The web application is open-source and available for use and modification under the MIT license.

